# Demonstration of a fast and easy sample-to-answer protocol for *Mycobacterium tuberculosis* diagnosis in point-of-care settings

**DOI:** 10.1101/601476

**Authors:** Nasir Ali, Graziele Lima Bello, Maria Lúcia Rosa Rossetti, Alexandre Dias Tavares Costa, Marco Aurelio Krieger

## Abstract

**Background:** Tuberculosis (TB) is currently the ninth leading cause of death worldwide and the leading cause from a single infectious agent, ranking above HIV/AIDS. Point of care diagnosis is one of the diagnostic aspects in the health care system that might have the potential to mitigate this worldwide epidemic. Although several qPCR tests are available, most cannot be taken to the field. Therefore, their use in POC settings is limited. Smooth sample preparation and streamlined DNA extraction constitute the biggest challenges for this limitation.

**Methods:** Seventeen *M. tuberculosis* samples which were already previously analyzed by GeneXpert or culture technique were subjected to our in-house protocol. Of these samples, ten were positive and seven negatives when tested by GeneXpert, while seven were positive and ten negatives when analyzed by culturing.

**Results:** Here we present a “proof of concept” protocol for sputum liquefaction and disinfection, followed by FTA card DNA extraction. The resulting DNA is rapidly amplified and *Mycobacterium tuberculosis* (MTB) DNA is detected with the use of a portable qPCR instrument. Our protocol is able to linearly identify down to 2 CFU/mL of MTB, showing great sensitivity on artificial samples. The protocol was challenged with patient samples, and showed excellent agreement with the gold-standard molecular protocol, allowing the detection of 9/10 positive samples (90%, or 10% of false negatives) and 7/7 of the negative (100%, no false positives). When compared to culture, 7/7 culture-positive samples were also found positive (100%, no false negatives), while 2/10 culture-negative were found positive by the present method (20% of false positives).

**Conclusion:** The proposed sample preparation protocol provides a rapid and easy procedure with a small number of reagents and steps, as well as minimal use of equipments, resulting in an easy-to-use tool for *M. tuberculosis* diagnosis in POC settings.

## Background

Tuberculosis (TB), caused by the bacillus *Mycobacterium tuberculosis* (MTB), is one of the leading causes of mortality from a single infectious microorganism, which disproportionately affects low and middle-income countries. In 2016, 1.7 million people have died and 10.4 million been fell ill due to TB, while the majority (56%) of the cases were from only five countries; India, Indonesia, China, Philippines and Pakistan (1). Although diagnostic tests based on amplification of nucleic acids (NAAT) such as the GeneXpert MTB/RIF have been widely used, a large gap of 4.1 million people without case notification is still posing a serious global health challenge.

The NAAT technology has open a new era of specific and sensitive diagnostic testing but these tests are mostly available to developed countries and resource rich areas (2). Sample preparation is the first step in molecular analysis and has got less attention among other steps like amplification and detection of genomic targets (3). Sample preparation has been a challenge for currently available molecular assays as well as a bottleneck in translating complex NAAT based molecular testing to easy-to-use point of care testing. Moreover, sputum samples for tuberculosis diagnosis pose additional challenges, since they are possibly a very highly contagious sample. The recommended procedure for tuberculosis diagnosis includes a step of sample inactivation using NALC/NaOH solution, where other bacteria are killed and the TB bacilli remains viable for culturing (4). For this purpose, a decontamination solution is prepared by adding of 4% NaOH and 0.5 g of NALC powder per 100 mL of the solution (5). This step is critical and is regularly performed together with sample liquefaction, thus rendering the sample more easily to handle.

Most of the molecular based assays developed to date, such as GeneXpert MTB/RIF depend on dedicated laboratory space and highly sophisticated equipment. A rapid, simple, efficient, less expensive and ideally equipment-free sample preparation would not only bring down the cost of molecular assays but also resolve the challenge of access to low-resource settings (6). Furthermore, the reagents used in molecular assays require special storage and transportation, which further increase the cost of the test (6). Largely, the successful integration of sample preparation with portable qPCR systems will solve the main hurdle in POCT diagnostic testing (4).The NAAT technology has open a new era of specific and sensitive diagnostic testing but these tests are mostly available to developed countries and resource rich areas (7)(8). Sample preparation is the first step in molecular analysis and has got less attention among other steps like amplification and detection of genomic targets (3). Sample preparation has been a challenge for currently available molecular assays as well as a bottleneck in translating complex NAAT based molecular testing to easy-to-use point of care testing. Most of the molecular based assays developed to date, such as GeneXpert MTB/RIF depend on dedicated laboratory space and highly sophisticated equipment. A rapid, simple, efficient, less expensive and ideally equipment-free sample preparation would not only bring down the cost of molecular assays but also resolve the challenge of access to low-resource settings (6). Furthermore, the reagents used in molecular assays require special storage and transportation, which further increase the cost of the test (6). Largely, the successful integration of sample preparation with portable qPCR systems will solve the main hurdle in POCT diagnostic testing (9)

Here we present a protocol for sputum samples that, coupled with FTA card extraction and integrated with a portable qPCR system, have the potential to translate the complex MTB molecular assay into a simple diagnostic platform.

## Methods

### Mucin solution 20 % (w/v) (1g/5mL)

A 20% mucin solution (M1778, Sigma Aldrich) was prepared in a 15 mL tube. After thorough mixing, the tube was placed in water bath for 2 h at 55 °C. Consistency of mucin solution was checked at intervals of 30 min with naked eye and gentle shaking to ensure proper solubilization. The tube was then placed for 4 h at room temperature (21-23 °C) and checked for the consistency at this point, the mucin solution, which at this point resembles a sputum sample. The solution was divided into 500 μL aliquots and placed at 2-8 °C in refrigerator for future use.

### Preparation of Solutions 1 and 2

Solution 1 was prepared by dissolving 7 g of guanidine thiocyanate (GSCN) powder in 10 mL of a 0.5 M EDTA solution (pH 8.0) in a 50 mL tube. The tube was heated at 65 °C for 1h. After the incubation, the volume was brought up to 25 mL with the same 0.5 M EDTA (pH 8.0) solution. The solution was divided into 500 μL aliquots and kept at room temperature for future use. Solution 2 consisted of commercial Tris-EDTA solution (Ambion™ Catalog Number AM9856).

### Patients and Clinical specimens

Seventeen sputum samples were received from Laboratório de Biologia Molecular (LBM) of the Universidade Luterana do Brasil (Canoas-RS, Brazil), which included 10 positive and 7 negative samples in three aliquots from individual patients. These samples were previously evaluated with GeneXpert MTB/RIF assay and culturing, so that the clinical history of the samples was already established. All the patients provided written informed consent (CAAE: 70697116.7.0000.5349).

### Sample preparation and application onto FTA card

One mL of the Sputum sample was mixed and thoroughly homogenized with 400 μL of Solution 1 in vortex for 20 to 30 seconds. Different volumes were also tested, but always keeping a ratio of 1.0 mL of sample to 0.4 mL of Solution 1. The homogenized mixture (sample plus Solution 1) was uniformly applied to an FTA Micro Elute card. The card was then allowed to dry for 1 hour at room temperature (21-23 °C).

### DNA extraction

A disc of 6 mm from the spotted area of the FTA card was punched with a sterilized puncher and placed in a clean 1.5 mL tube. Five hundred microliters of Solution 2 were added to the tube and vortexed for 3 times of 5 seconds with 5-second intervals. Next, the tube was incubated at 95 °C for 5 minutes. The tube was cooled down to 25 °C and the supernatant was collected into a new tube for further use. The supernatant contained the extracted DNA that was further used for qPCR analyses. The remaining card was stored at room temperature for future uses.

### Analytical sensitivity

To evaluate the analytical sensitivity of the DNA extraction technique on FTA card, an 8-point serial dilution was performed from the reference strain H37Rv of *M. tuberculosis*. Subsequently, individual mucin samples were each contaminated with a dilution (25 μL in volume) and were equally distributed on individual cards. DNA extraction was performed from each of these eight samples, and the portable qPCR (Q3-Plus) was used to evaluate the detection limit of the method described herein. For the counting of CFUs, each of the 8-point serial dilution was inoculated (25 μL in volume) in petri plates (in duplicate) containing culture medium for mycobacteria growth (7H10 agar).

### Experimental conditions for regular qPCR

For detection of MTB, primers and probe for the IS6110 gene were purchased as a custom Taqman® gene expression assay (Applied Biosystems, catalog number P170804-006F09-250917). The reaction was performed on the standard instrument ABI7500 (Applied Biosystems) or on a portable prototype instrument named “Q3-Plus” developed by STMicroelectronics (Agrate Brianza, Italy)(10). The Q3-Plus system has small dimensions (7 × 14 × 8.5 cm), weighs only 150 grams, performs the thermal cycles in a disposable silicon chip (Figure1), and can be operated with any small portable computer. For the ABI7500 reaction, the reaction contained the Multiplex PCR Master (IBMP, Curitiba, Brazil) and MTB Taqman Oligomix (which includes primers and probe; assay name IS6110-A, assay ID: AIPACNK, Thermo Scientific, USA) was used. Reactions were standardized to the final volume of 20 μL, considering the addition of 5μL of sample (extracted DNA). In the ABI7500 instrument, the cycling conditions were 50 °C for 2 minutes, 95 °C for 10 minutes, followed by 45 cycles of 95 °C for 15 seconds and 60 °C for 60 seconds. For the Q3-Plus reaction, one microliter of the extracted DNA was mixed with 0.25 μL of MTB Taqman Oligomix, and 1.75 μL of Multiplex PCR Master (IBMP, Curitiba, Brazil) in a final volume of 5 μL. For the Q3-Plus system, the cycling conditions were: 80 °C for 10 seconds, 97 °C for 60 seconds, followed by 45 cycles of 97°C for 20 seconds and 62 °C for 60 seconds; and optical parameters for the FAM channel were exposure time of 1 second, led power of 3 and analog gain of 15, while for the VIC channel the optical parameters were exposure time of 2 seconds, led power of 8, and analog gain of 14. Reactions on both instruments were also supplemented with oligonucleotides for the detection of the human 18SrRNA gene (Table 1).

**Figure 1.**
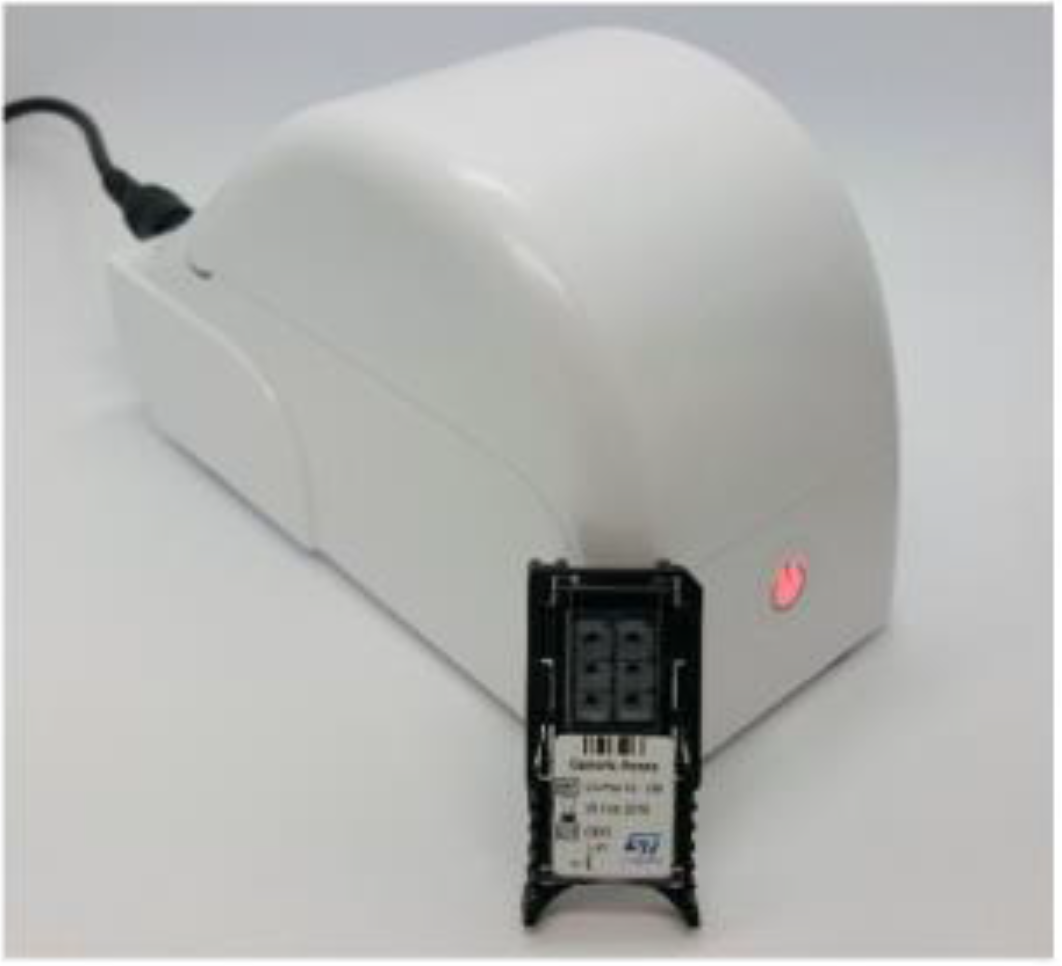
Q3-Plus platform with its silicon chip. The Q3-Plus instrument with the lid closed, and the disposable unit (silicon-chip cartridge)

**Table 1.**
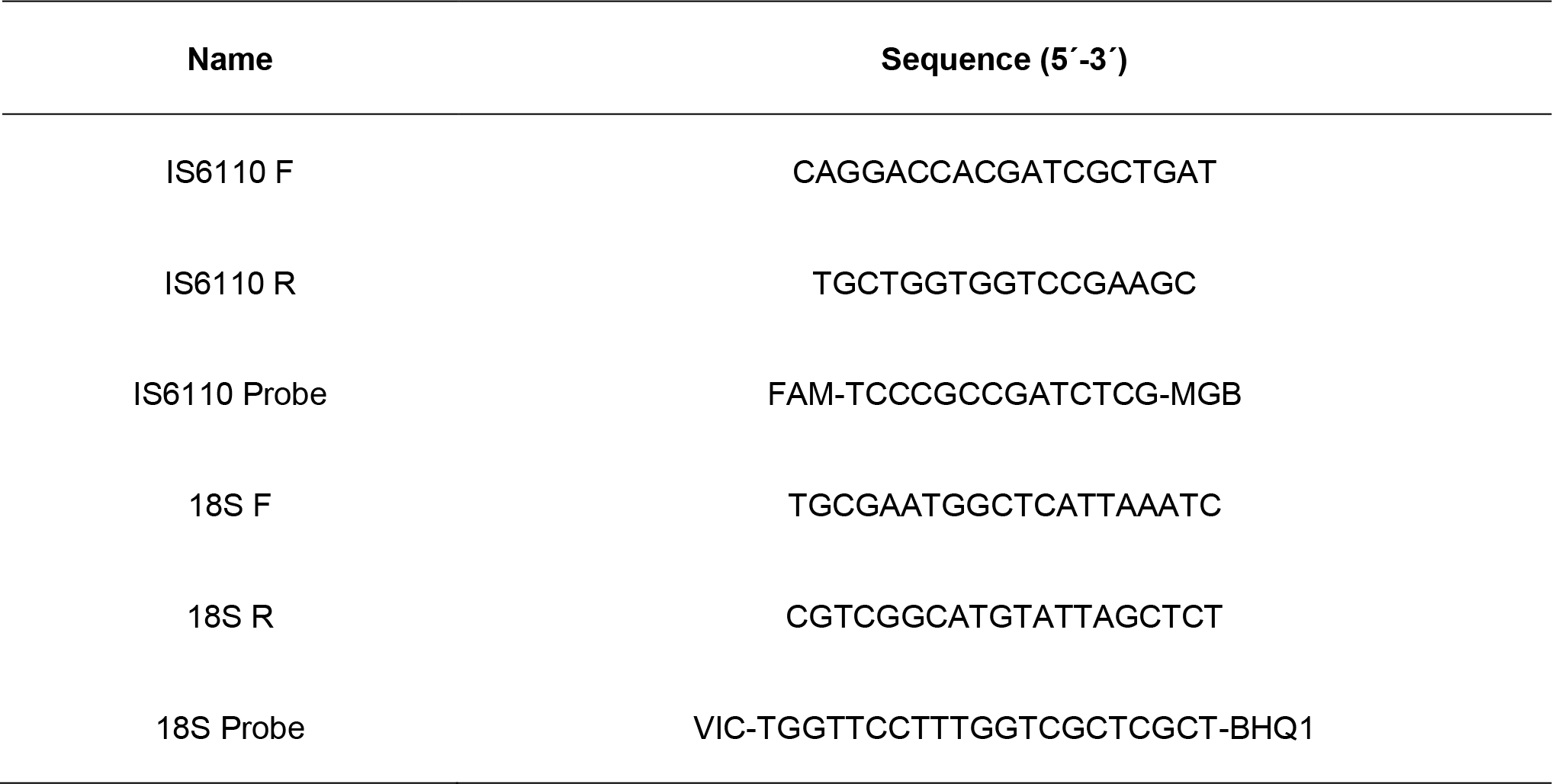
Sequence of primers and probes used for the target IS6110 of MTB and Internal control

### Gelification of qPCR reagents

Gelification of the qPCR reagents was performed by adding the Gelification solution in substitution for water in all qPCR reactions, according to published protocols (11)(12)(13). The Gelification solution contains three classes of components, each with specific functions within the mixture and the process: (i) compounds that protect the biomolecules from the desiccation process by stabilizing and reducing the activity of water in the solution; (ii) free radical scavengers, which inhibit oxidizing reactions between carbonyl or carboxyl groups and the biomolecule’s amino or phosphate groups; and (iii) inert polymers, which creates a broader protective matrix. These components define a sol-gel mixture that prevent the loss of the tertiary or quaternary structure during the desiccation process, maintain the biomolecules’ activity upon rehydration. The qPCR mix containing the gelification solution and all qPCR reagents (enzymes, buffers, oligonucleotides, dNTPs, etc.) was manually aliquoted to each reaction well, and the reaction vessels (either the ABI7500 plastic 96-well plates or the Q3-Plus silicon chips) were submitted to vacuum (30 ± 5 mbar) under controlled temperature (30 ± 1 °C) to gelify the reagents. For all experiments, gelified reactions were stored at 2-8 °C for up to 7 days. The same threshold parameters and positivity criteria were used for detection of MTB or human genomic targets.

## Results

The first step we tackled was the initial inactivation and liquefaction of the sputum sample. Since we would like our procedure to be used in the field or in low-resource settings, we were not considering culturing as a viable diagnosis method. Therefore, we envisioned that the inactivation of the sample could as well inactivate all bacteria, TB bacilli included.

In a previous study, Triton X-100, urea and GSCN were shown to kill all bacteria in sputum samples, including MTB (4)(14). Therefore, we evaluated if concentrated GSCN would be able to liquefy an artificial sample consisting of pig mucin, which can be used as a substitute for sputum (15). GSCN at 6M was used alone, or together with other chaotropic and dissolving chemicals, such as urea (8 M) or Triton X-100 (at 10%). Similar results were obtained if SDS (at 10%) substituted for Triton X-100.

GSCN, urea and Triton X-100 been experimented with different volumes and combinations. A sample volume of 500 μL of porcine mucin was chosen, as it seems realistic volume to obtain from a patient, and the consistency is similar to human sputum. In the context of sample preparation, NALC/NaOH was added to the standard sample in same volume i.e. 500 μL. The initial viscosity of porcine mucin was rated as (++++), and after treatment with chemical(s) it was rated from ++++ (most viscous) to + (least viscous). Viscosity was observed and measured with the naked eye while gently agitating the tubes. The results obtained are shown in Table 2. Tube 1 was considered as “control sample” and prepared according to the protocol used by WHO with some modifications i.e. not using phosphate buffer solution. Triton X-100 alone was the least effective component in the liquefaction process, while urea alone or in combination with Triton X-100 showed similar results (++), without any profound effect on liquefaction. Addition of GSCN alone was very effective (+), while its combination with urea or Triton X-100 did not further decrease the viscosity. Taken together, these results suggest that Urea contributes to the reduction of viscosity, but the most effective adjuvant was GSCN. It is important to emphasize that NALC/NaOH was not added because this sample will not be used for culturing.

**Table 2.**
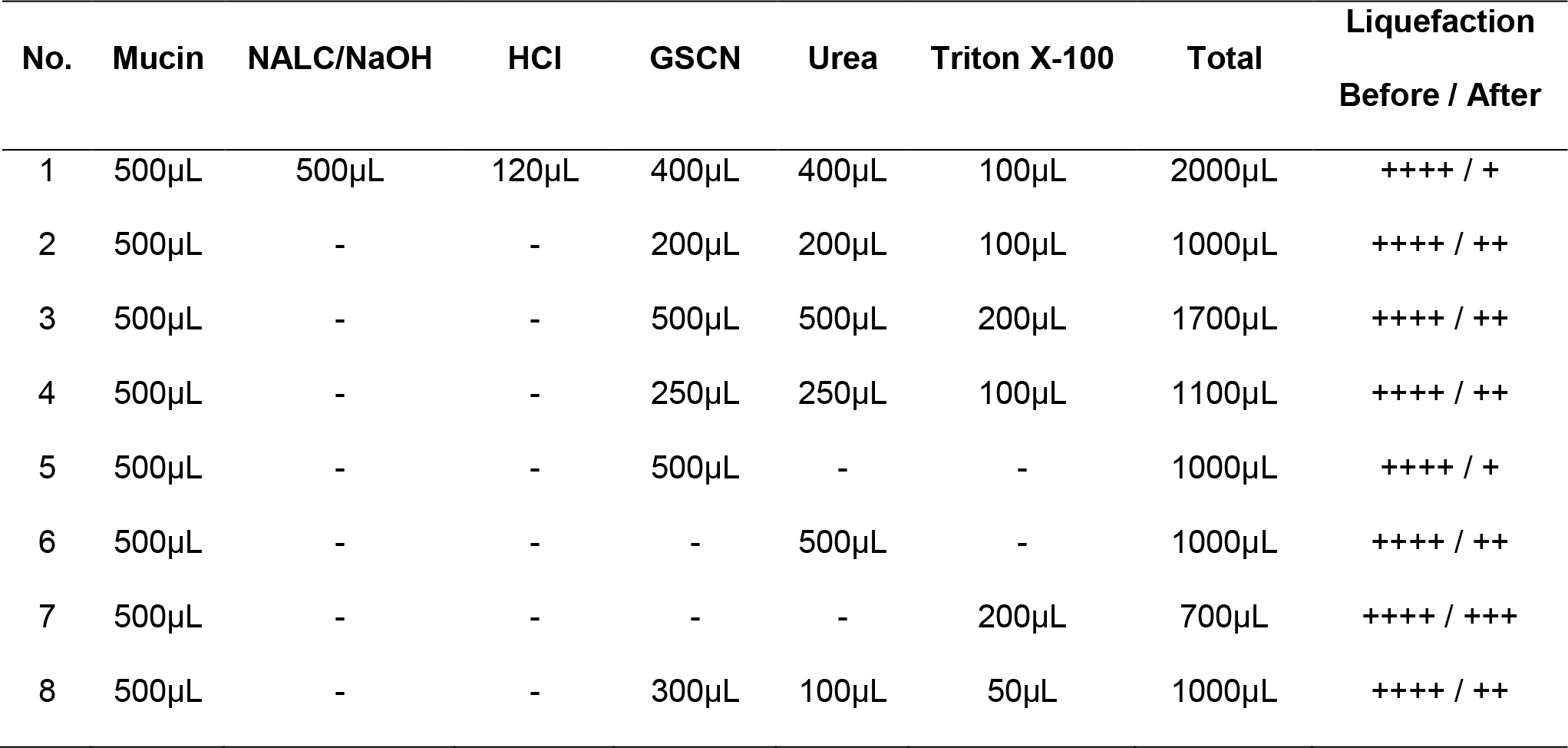
Liquefaction of mucin with different chemicals

Next, we evaluated different extraction protocols mentioned in Table 3. We used six protocol variants based on different punch size (1 punch, 3 punches or whole circle), washing and elution buffers (H_2_O or TE pH8), as well as different incubation conditions (with and without incubation). For this purpose, 500 μL of mucin was selected as sample size and spiked with 25 ng/μL of DNA from H37Rv cells.

**Table 3.**
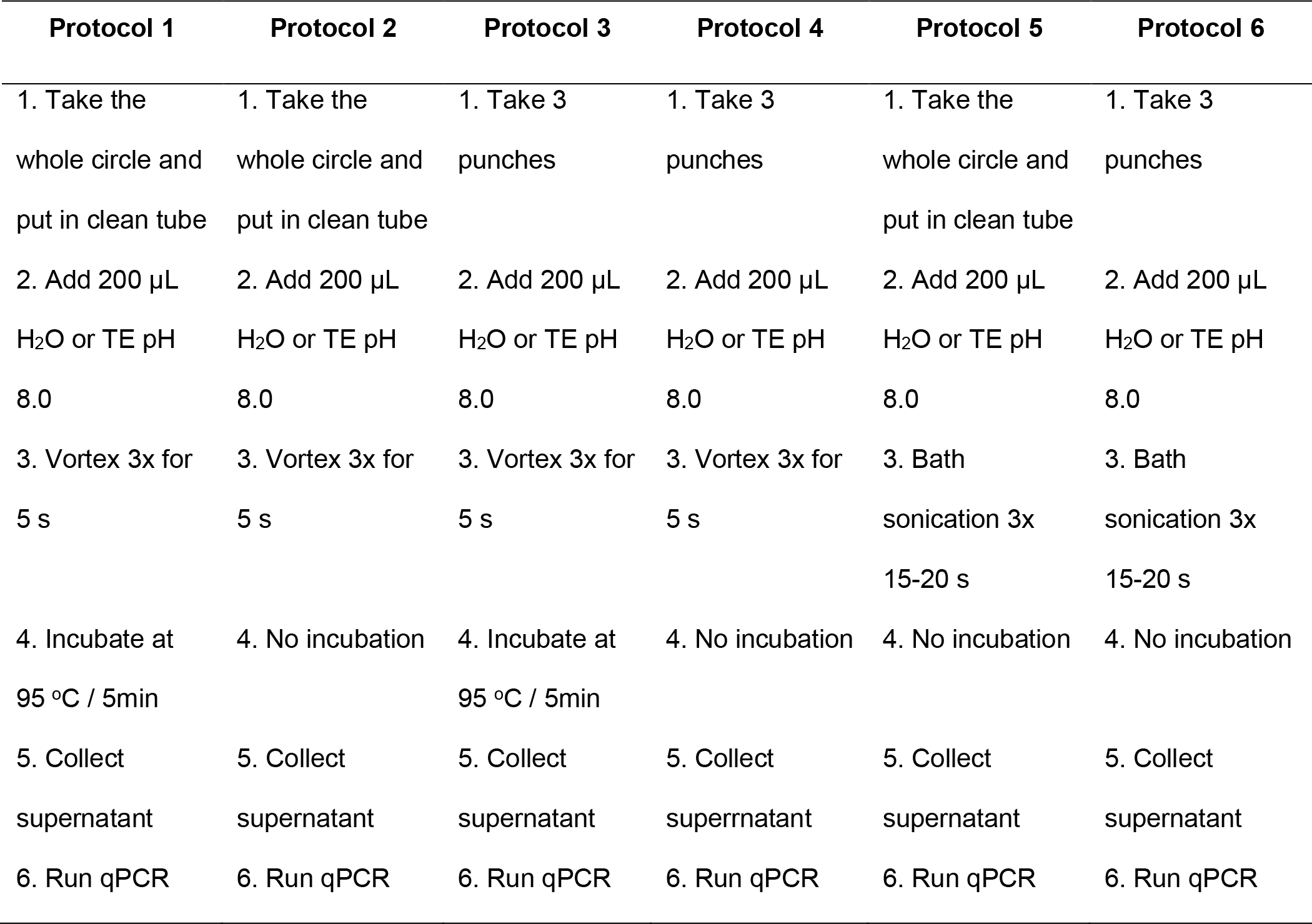
Six different DNA extraction protocols

At the end, protocols 3 and 4 proved to be better as compared to the others (1, 2, 5, 6). Protocol 3 and 4 (having 3 punches) were different from protocol 1 and 2 (having whole circle) with respect to punch size, while 5 and 6 different in incubation conditions (sonication). As a lysis buffer, 350 μL of GSCN was added and extraction was done for all the protocols, but only 3 and 4 showed amplification (Figure 2).

**Figure 2.**
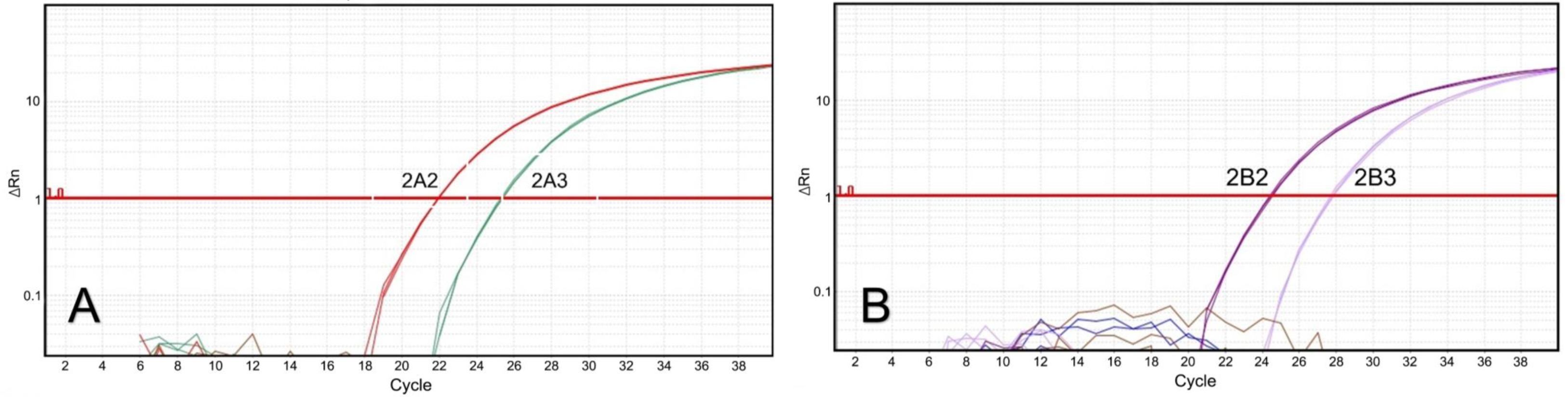
Real time PCR representative traces obtained with DNA extraction protocols 3 or 4. Panel A shows the detection of *M. tuberculosis* DNA using Protocol 3 (three punches of 6 mm in FTA card, with incubation at 95 °C). Panel B shows the detection of *M. tuberculosis* DNA using Protocol 4 (three punches of 6 mm in FTA card, without incubation at 95 °C). In both cases, there is no amplification with the undiluted DNA extract, but there are positive amplifications when the extract was diluted 1:10 and 1:100, suggesting presence of inhibitors in the undiluted DNA extract.

**Figure 3.**
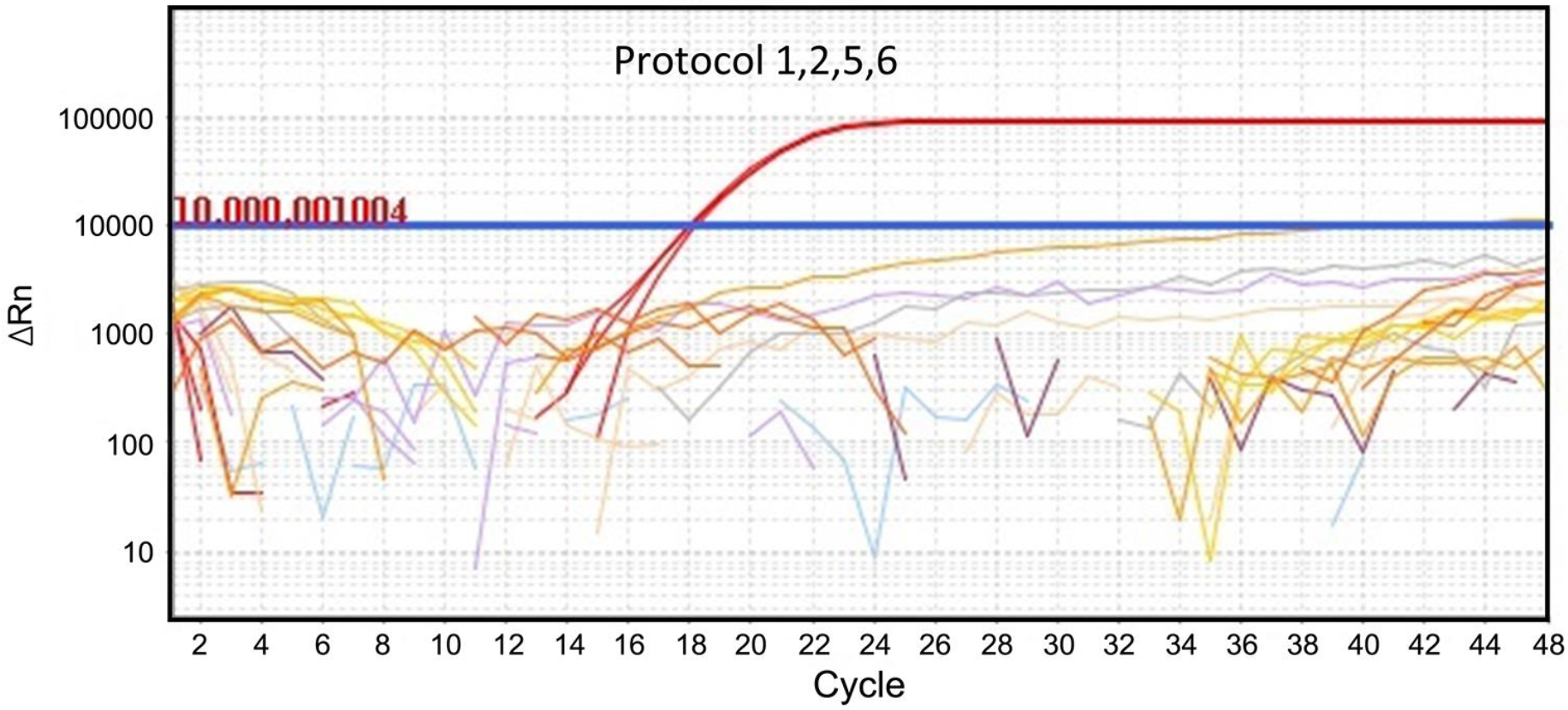
DNA extraction with other protocols yielded no detectable amplification. Four extraction protocols did not show any detectable amplification of *M. tuberculosis* DNA, even when diluted at 1:10 or 1:100 ratios, while detection of the human endogenous control was possible in the diluted samples (red lines).

Interestingly, the direct extract did not show any amplification while diluting both the samples further for 1:10 and 1:100 showed positive amplification, suggesting that some contaminants and inhibitors are present in the extract (Figure 2). Table 4 summarizes the Ct values obtained for each dilution in protocols 3 and 4.

**Table 4.**
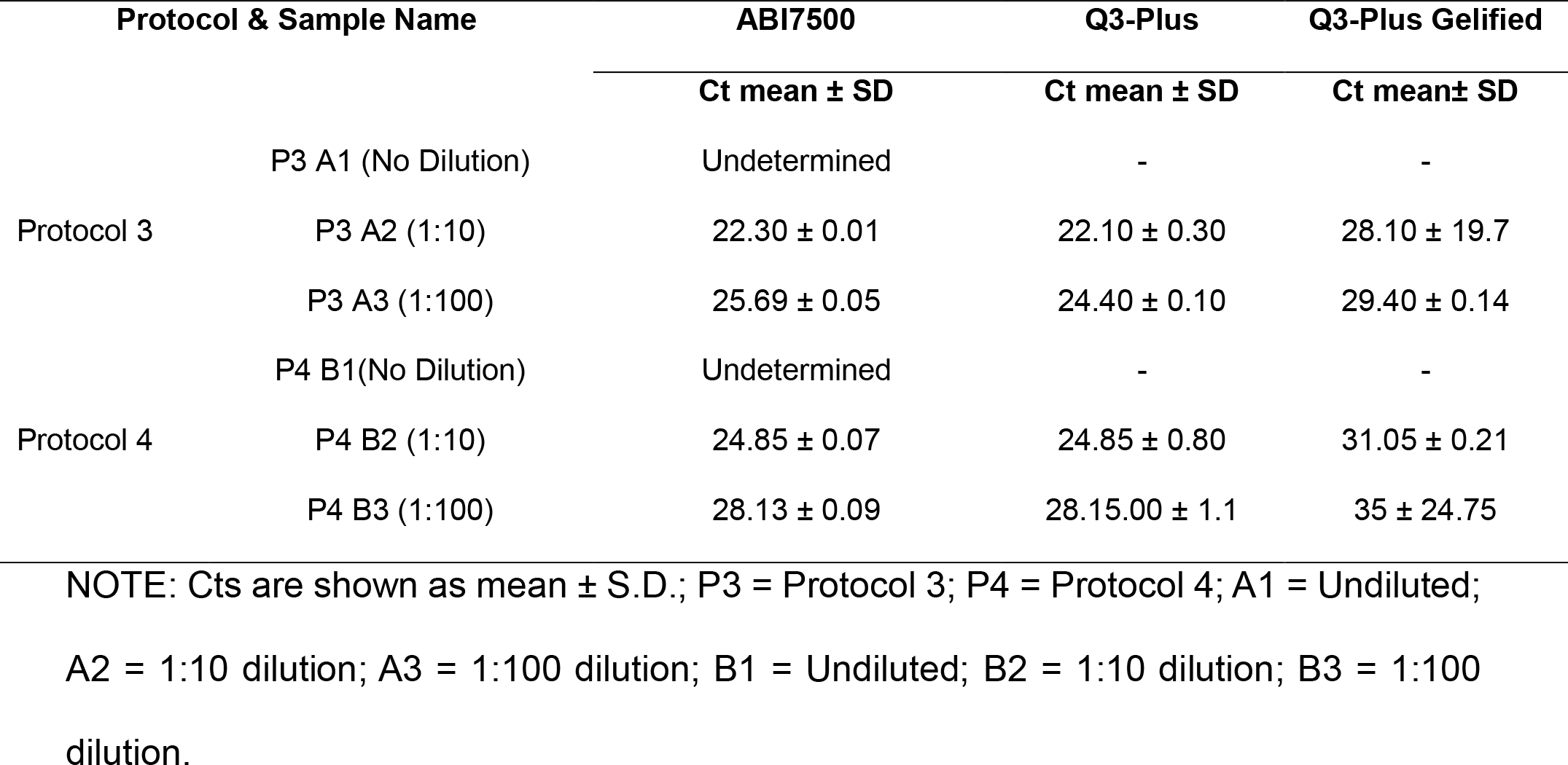
Comparison of ABI7500, Q3 with Protocol 3 and Protocol 4.

### Evaluation of FTA extraction protocol with whole MTB cells and CFU count

In order to evaluate our in-house protocol for whole MTB cells, a 7-point dilution of whole MTB cells (strain H37Rv) were spiked and applied on FTA cards. Following extraction, real time and Q3 PCR was performed. On the other hand, all the dilution points were inoculated on culture plates for CFU count. By evaluating the analytical sensitivity, the detection limit of DNA extraction from the FTA card detected DNA from the microorganism on the Q3-Plus until the dilution of up to 1:4 and the average corresponding was two CFU count.

### A comparative study of different techniques for MTB Samples

All the 17 MTB samples were tested previously by GeneXpert PCR and culture. The untreated samples were submitted to the simplified In-house protocol. The extracted DNA was then evaluated on the standard (ABI7500) and the portable (Q3) instruments. The data collected for all samples are mentioned in Table 5. As can be seen from the table the samples tested by our protocol showed similar results as compared to GeneXpert and the gold standard culture technique. Overall, these results suggest that our in-house protocol has the potential to be used in future for detection of MTB. Table 5 shows that our in-house protocol showed almost similar results as GeneXpert, only one sample gave a different result (sample #10). All other, negative and positives, gave the same results. Not surprisingly, when GeneXpert results differ from culture, ours differed too.

**Table 5.**
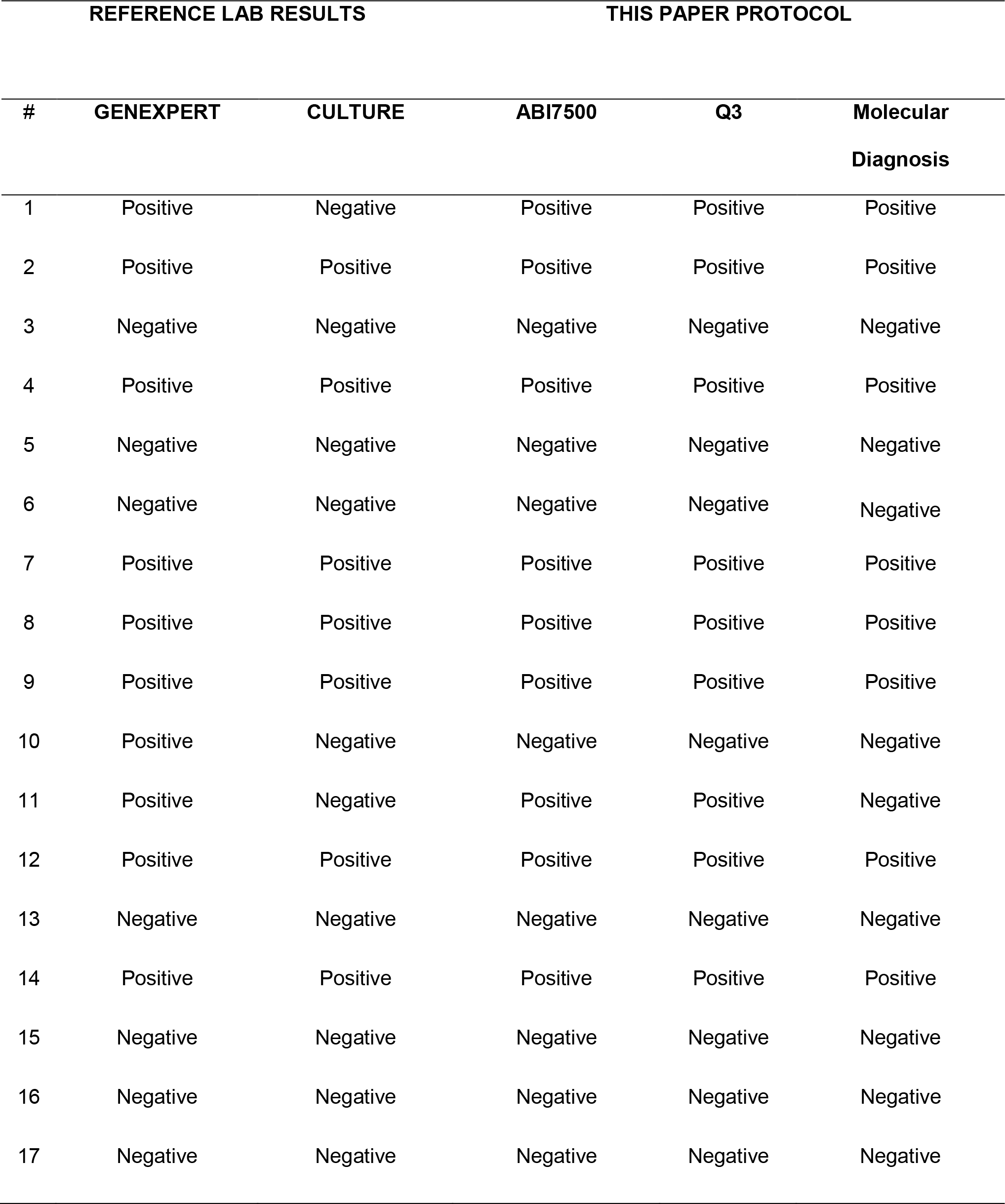
Comparison among different platforms for detection of MTB.

## Discussion

In this study we present an optimized and streamlined assay for the diagnosis of MTB using a portable qPCR system (16). Tuberculosis is the leading infectious cause of death ranking above HIV/AIDS and ninth in most lethal disease worldwide. The disease disproportionately affects low and middle-income countries due to weak health infrastructure. This study set out with the aim of finding a diagnostic methodology for *M. tuberculosis* which match the WHO ASSURRED criteria: <u>a</u>ccurate, <u>s</u>ensitive, <u>s</u>pecific, <u>u</u>ser friendly, <u>r</u>apid, <u>r</u>obust, <u>e</u>quipment-free and <u>d</u>eliverable to end user (1). All these requirements have been met (if not all) to a larger extent in different platforms such as GeneXpert Ultra and Bio Fire Diagnostics, our Q3-Plus showed some advantages over the standard ABI7500 in terms of equipment cost, reaction volume, user friendly interface, fast temperature ramping, shorter total reaction time

Smooth sample preparation and streamlining of DNA extraction to an efficient detection method is one of the challenges for currently available molecular diagnoses of *M. tuberculosis*(17), which results in their limited use in resource-constrained areas. In this study, our aim was to demonstrate a rapid, simple, and cost-effective sample preparation and DNA extraction protocol for *M. tuberculosis*, integrating paper-based DNA extraction and specific target amplification/detection by a portable and easy-to-use real time PCR platform. The final experimental overview of the fully developed procedure is outlined in Figure 4.

**Figure 4.**
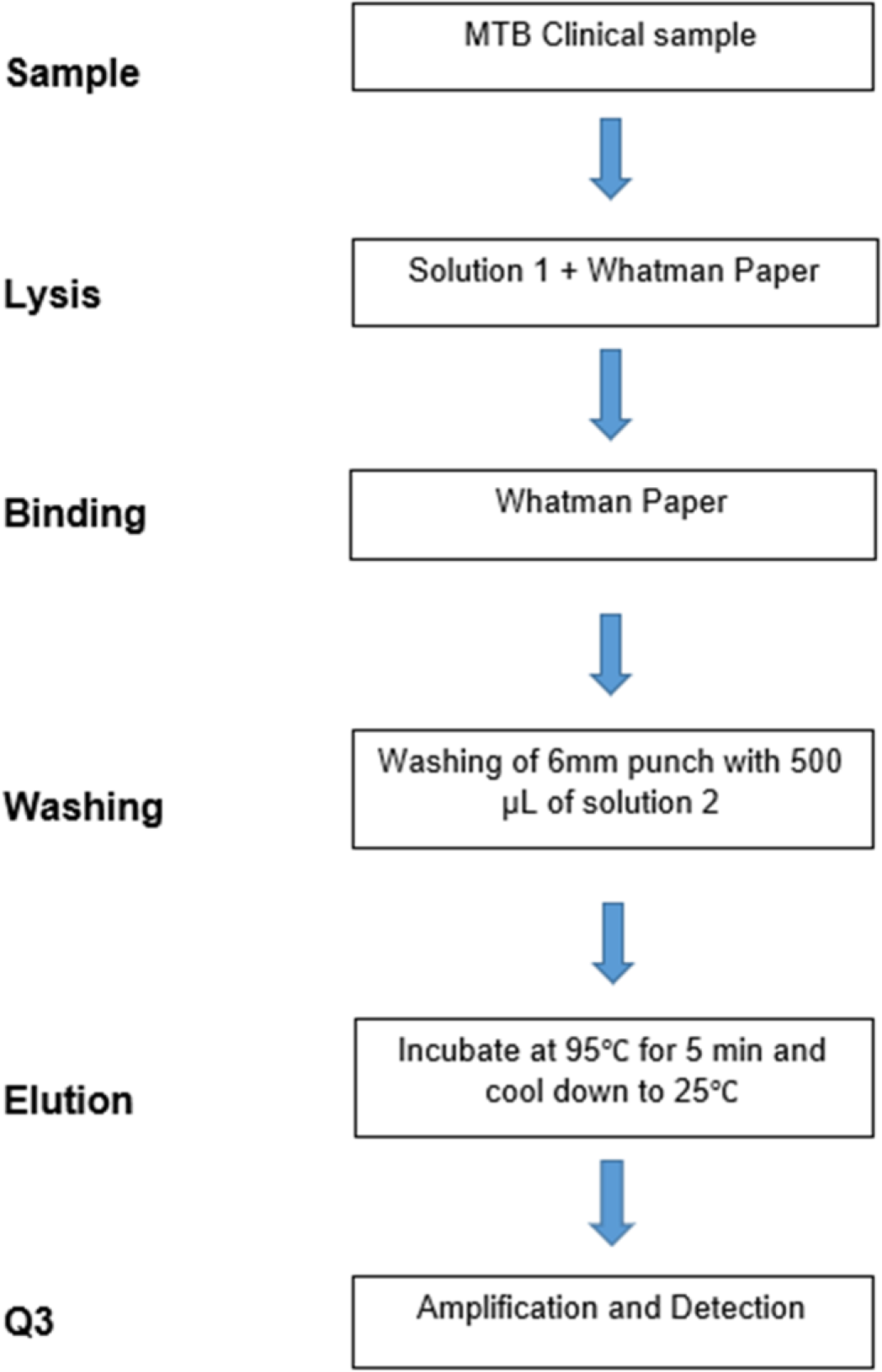
Overview of the proposed simplified and streamlined method. Flow chart depicting the different steps of the present study’s DNA extraction protocol, comparing with the steps of traditional methods.

To disrupt the complex network of interlinked mucin matrix, which is the main culprit for trapping the target microorganism, a mechanism is needed to liquefy the sputum thus releasing the pathogen(5). Different strategies (chemical, mechanical or combination of both) can be applied for liquefaction of the sputum sample, in a process where it changes from viscous heterogeneous to a homogenous/solubilized liquid state (17). In the current study, several chemicals were screened for homogenization of sputum sample such as the chaotropic agents GSCN and urea, or the detergents NP-40, SDS, and Triton-X 100. Interestingly, GSCN by itself showed very efficient results in terms of liquefaction, even when combined with the other tested compounds. In a previous study, PAWLOWSKI and KARALUS (18) have shown that highly GSCN can be used in sample preparation for the extraction of DNA and protein for a variety of samples (including liquid samples and in case solid samples such as food and stool must be suspended in PBS or water). More importantly, *M. tuberculosis* was shown to become non-viable after exposure to commercial lysis buffers, in which GSCN is one of the main components (4). This GSCN’s feature is essential to the protocol developed in the present work, since we propose the sample to be manipulated in low-resource settings, which might not have safety cabinets.

FTA elute card™, based on Whatman FTA technology (Whatman, Inc.), are chemically treated papers that enable cell lysis upon contact and release nucleic acids (19). Integrating liquefaction and decontamination of the sputum sample to FTA-based extraction is another approach for TB diagnosis adopted in this protocol. Here, we applied elution approach for MTB in sputum samples tested with different size of punches and elution buffers modifying the manufacturer’s protocol. GOVINDARAJAN et al. (15) have demonstrated a rapid and cost-effective sample preparation using a microfluidic “origami”, using spiked mucin with *E.coli* as a model. As compared to the microfluidic “origami”, the FTA card is more efficient in cell lysis and disintegration of the viscous matrix of a sputum sample because of the dry reagents already stored on the paper. Moreover, microfluidic origami which takes around 2 h while our protocol takes 1h 30min from raw sample to DNA extraction with need solely for a heat block and with minimal use of liquid reagents. The use of FTA cards for storage, transport and preservation of sputum samples has been performed successfully for diagnosis of MTB by (20), who have shown that MTB DNA was stabilized with FTA card for six months at room temperature. These findings provide not only a simple and economically favorable solution for collection, storage and transportation of the sputum but also suggest the use of these cards for further sample preparation and extraction.

Both GeneXpert and culture are considered as reliable MTB diagnostic assays for sputum samples. However, these techniques present challenges for application in POC-Dx. When compared to our in-house protocol with the early results obtained with GeneXpert and culture, our protocol shared almost same results with both techniques. Although the low number of samples analyzed in this study is a limiting factor, these results are comparable to gold standard techniques in case of detection with the hands-on time of around 3h and can be applied in the future as a triage test in resource-limited areas. In addition, our results are in accordance with the culture and GeneXpert results. Thus, it has the potential to be used in high TB burden countries such as India, Indonesia, China, Philippines and Pakistan, after testing with more characterized samples not only with Q3 liquid reaction but also in a gelified format as well. The gelified reactions performed similarly to conventional reactions and could be implemented in laboratories and small medical centres. The insertion of an internal control of the reaction, the human 18S rRNA, guarantees us the functionality of the system as a whole, with respect to sample extraction, nucleic acid integrity, the absence of inhibitors in the reaction, and correct functioning of equipment and software.

For comparison purposes, the cost of DNA extraction with a kit from Roche is US$ 5 per sample, while the cost for our extraction protocol using GSCN+FTA paper can be roughly estimated at US$ 1-2 per sample. The cost comparison of the PCR reaction per patient between a Q3-plus instrument and ABI 7500 is almost the same (US$3-4) while there is a huge difference in cost of the instrument (Q3-plus < US$10,000 while ABI7500 > US$40,000) (21)(22). In general, ABI7500 needs skilled professionals, sophisticated lab environment and requires time specific maintenance and calibration while Q3-Plus platform relies on minimum personal training may be one day and does not require maintenance/calibration or other lab infrastructure.

Turnaround time is an important aspect of any diagnostic assay. The culture technique, which is the gold standard for MTB diagnosis, can take 4 to 16 weeks (23), and such a long period can result in further complexities such as delays in initiation of the treatment or loss of disease follow up. On the other hand, GeneXpert Ultra have turnaround time of 80-90 min but have relatively high cost (concessional price is 9.98 US$ for eligible countries). We have estimated the turnaround time for our tests as 3 hours from raw sample to result in assay. This feature provides a rapid diagnosis that, coupled with the accurate molecular detection elicited by qPCR, leads to an early and more efficient start of treatment.

Our assay reagents are workable at room temperature without the need for any special storage infrastructure, which makes tuberculosis diagnosis feasible in poorly equipped areas. Furthermore, access to quality diagnosis at POC will not only allow timely initiation of treatment but also reduce the dropout rate of the patients, which is one of the key factors for the spread of TB disease (24).

## Conclusions

The present work shows the development of a simple and inexpensive sample preparation protocol for the detection of TB in viscous sputum sample, integrated with an easy-to-use portable real time PCR system, thus comprising a simple sample-to-answer diagnostic tool. Our sample preparation protocol allows the detection of an MTB bacterial load as low as two CFU. The work presented here is a step towards point of care diagnosis for MTB, being suitable for low infrastructure primary health units as well as secondary care health clinics.

## Acknowledgements

N.A holds CAPES fellowship at Departamento de Engenharia de Bioprocessos e Biotecnologia, Universidade Federal do Paraná (UFPR). This work was partially funded by a grant from the Banco Nacional de Desenvolvimento Econômico e Social (BNDES), contract no. 15.2.0473.1 (Operation #4.816.864). CAPES, UFPR, or BNDES had no participation in the study’s design, data collection, analysis, interpretation, or writing of the report, and decision to submit for publication.

